# *Drosophila* architectural protein CTCF is not essential for fly survival and is able to function independently of CP190

**DOI:** 10.1101/2021.05.31.446447

**Authors:** Olga Kyrchanova, Natalia Klimenko, Nikolay Postika, Artem Bonchuk, Nicolay Zolotarev, Oksana Maksimenko, Pavel Georgiev

## Abstract

CTCF is the most likely ancestor of proteins that contain large clusters of C2H2 zinc finger domains (C2H2) and is conserved among most bilateral organisms. In mammals, CTCF functions as the main architectural protein involved in the organization of topology-associated domains (TADs). In vertebrates and *Drosophila*, CTCF is involved in the regulation of homeotic genes. Previously, null mutations in the *dCTCF* gene were found to result in death during the stage of pharate adults, which failed to eclose from their pupal case. Here, we obtained several new null *dCTCF* mutations and found that the complete inactivation of dCTCF appears to be limited to phenotypic manifestations of the *Abd-B* gene and fertility of adult flies. Many modifiers that are not associated with an independent phenotypic manifestation can significantly enhance the expressivity of the null *dCTCF* mutations, indicating that other architectural proteins are able to functionally compensate for dCTCF inactivation in *Drosophila*. We also mapped the 715–735 aa region of dCTCF as being essential for the interaction with the BTB (Broad-Complex, Tramtrack, and Bric a brac) and microtubule-targeting (M) domains of the CP190 protein, which binds to many architectural proteins. However, the mutational analysis showed that the interaction with CP190 was not essential for the functional activity of dCTCF *in vivo*.

**Highlights:**

- The dCTCF 715–735 aa region interacts with the BTB and M domains of CP190
- Interaction with CP190 is not essential for dCTCF function *in vivo*
- Null *dCTCF* mutants are viable but show severely reduced fertility

## 1. Introduction

The chromosomes of higher eukaryotes are organized into a series of discrete, topologically independent regions known as topology-associated domains (TADs) [1-3]. Dynamic interactions between enhancers, silencers, insulators, and promoters occur predominantly within these domains [4-7]. Recent studies in single cells have shown that domain boundaries only determine the greatest likelihood of interactions between regulatory elements within the TAD without completely blocking interactions between regulatory elements located in neighboring TADs [2, 8].

In *Drosophila*, several proteins that contain arrays of C2H2 zinc finger (C2H2) domains display architectural and insulator functions [9, 10]. One architectural protein, CTCF, is conserved among most bilateral organisms and is the likely ancestor of all C2H2 proteins [11, 12]. CTCF contains a cluster of 11 C2H2 domains localized in the central part of the protein. The five most conserved C2H2 domains recognize a specific 15-bp consensus site that is commonly found in all studied organisms [12, 13]. The N-terminal regions of CTCFs contain unstructured domains that are able to homodimerize [14]. In mammals, CTCF has been shown to be a key architectural protein that supports distance interactions between enhancers and promoters [15-17] and is involved in the formation of TAD borders [16, 18-20]. According to a prevalent model, CTCF-binding sites block the movement of cohesin complexes along DNA, resulting in the formation of chromatin loops [21]. Recently, the evolutionary conserved 222–231 aa region of the human CTCF (hCTCF) protein was found to interact with cohesin and appears to be essential for chromatin loop formation [22, 23].

CTCF sites are commonly found in mammalian genomes, and CTCF inactivation typically results in cell death and embryonic mortality, in addition to significantly disturbing the chromosomal architecture [16, 24, 25]. However, the inactivation of hCTCF only affects the expression of a small number of genes [16], which is consistent with a model in which CTCF is the primary but not the only architectural protein in mammals.

*Drosophila* CTCF (dCTCF) binds predominantly to gene promoters, introns, and intergenic regions [26-28]. The most evident functional role for dCTCF involves the organization of boundaries between regulatory domains in the *bithorax* complex (BX-C). BX-C (Fig. 1A) consists of three highly conserved homeosis genes, *Ubx, abd-A*, and *Abd-B*, which encode transcription factors that determine the identity of the last thoracic and all abdominal segments of the fly [29]. Mutations in the *dCTCF* gene predominantly affect the proper expression of *Abd-B*, which is responsible for the development of abdominal segment 5–9 (A5–A9) [30-33]. The deletion of specific CTCF-binding sites or large regions containing CTCF-binding sites leads to the promiscuous activation of *Hox* developmental genes in mammals [15, 34, 35], suggesting an evolutionarily conserved role for CTCF in the regulation of homeotic genes.

**Figure 1.**
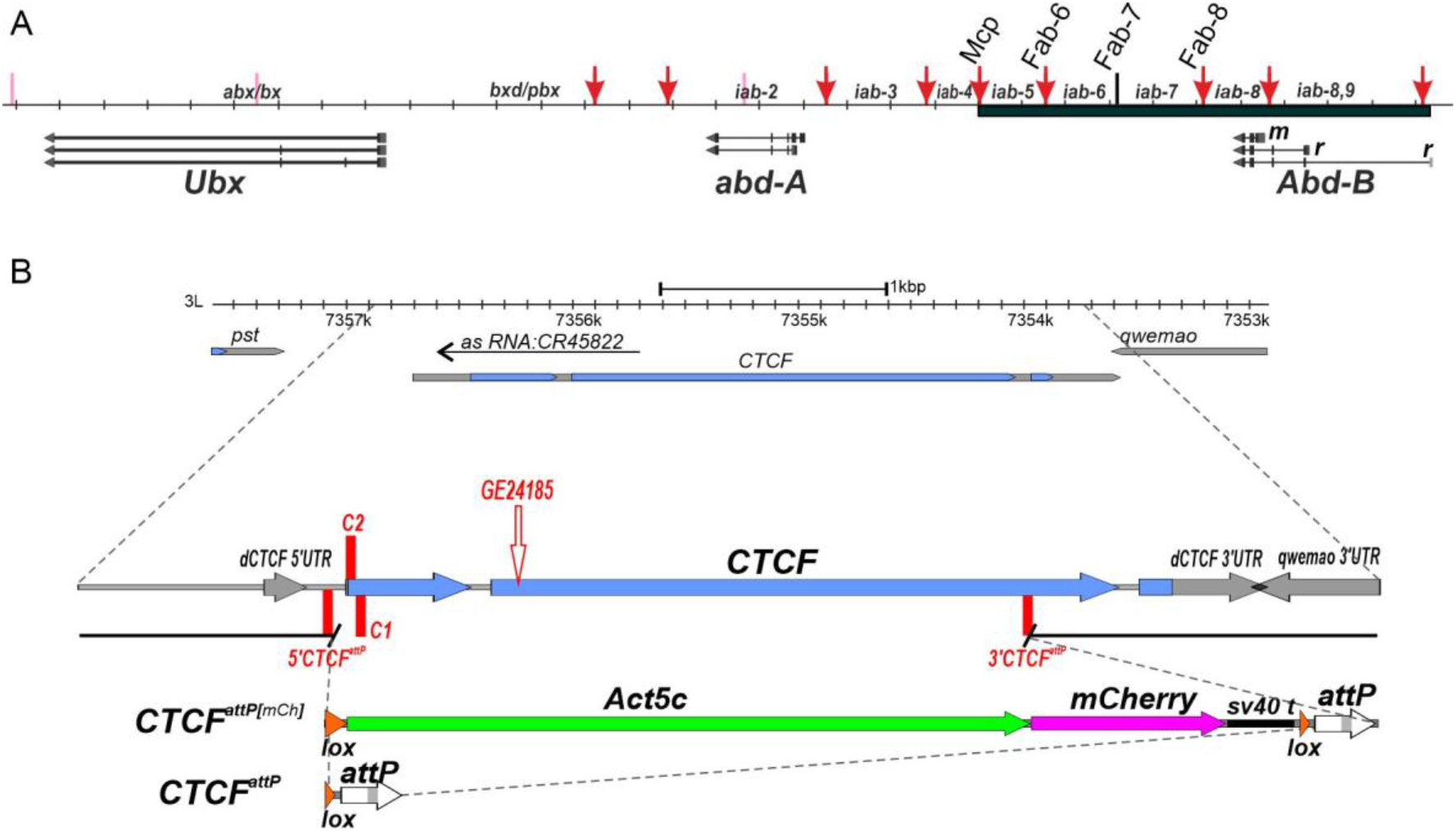
(A) Schematic diagram of *BX-C* (presented as a sequence coordinate line). The *Ubx, abd-A*, and *Abd-B* transcripts are marked by horizontal black arrows. The mapped dCTCF-binding sites in the boundaries are indicated above the coordinate map as red vertical arrows. The dCTCF-binding sites are located in the *Mcp, Fab-6*, and *Fab-8* boundaries and in the proximal (m) and distal (r) promoter regions of *Abd-B*. The expression of *Abd-B* in the A8–10 segments is regulated by the distal promoter. In the *Abd-B* regulatory region, only the *Fab-7* boundary (vertical black bar) does not contain any dCTCF-binding sites. (B) CRISPR/Cas9 editing of the *dCTCF* gene. The *dCTCF*-coding regions are shown as blue horizontal arrows. The surrounding genes (*pst* and *qwemao*) and the untranslated regions of the *CTCF* gene are shown as gray arrows. The insertion site for the transgene in the *GE24185* mutation is indicated by the empty red arrow. CRISPR targets are shown as vertical red bars. The proximal and distal endpoints of the *CTCF*^*attP[mCh]*^ deletion are indicated by breaks in the black line. The *mCherry* reporter (magenta arrow), controlled by the *Act5C* promoter (green arrow), was used for the selection of the *dCTCF* deletions. The *attP* and *lox* sites were used in genome manipulations and are shown as white and orange arrows, respectively.

The dCTCF protein contains a folded N-terminal domain that forms tetramers in solution [14]. The deletion of the N-terminal domain from dCTCF affects all *in vivo* functions of the protein, including the regulation of *Abd-B* expression [36]. The primary binding partner of dCTCF is the CP190 protein, which belongs to a large family of transcription factors containing an N-terminal BTB (Broad-Complex, Tramtrack, and Bric a brac) domain [30, 36, 37]. Several studies have identified multiple DNA-binding proteins that are able to recruit CP190 [38-44]. The available data have suggested a role for CP190 in the formation of open chromatin structures and the recruitment of the nucleosome remodeling factor (NURF), dimerization partner, RB-like, E2F, and multi-vulval class B (dREAM), and Spt-Ada-Gcn5 acetyltransferase (SAGA) complexes [27, 45-52].

To further study the functional roles of dCTCF *in vivo*, here we generated two null point mutations and deleted the *dCTCF* gene using a CRISPR/Cas9-dependent genome editing technology [53, 54]. As previously demonstrated, these *dCTCF* mutations primarily affected the expression of the *Abd-B* gene. We also found that different genomic backgrounds affected the expressivity of the null *dCTCF* mutations by increasing the lethality and sterility of flies. Co-immunoprecipitation and *in vitro* pulldown experiments showed that the 705–733 aa region of dCTCF is critical for mediating the interaction with the CP190 protein. The expression of a mutant dCTCF protein that lacked the CP190 interacting domain (dCTCF^Δ705–733^) was able to completely compensate for the *dCTCF* null mutation, suggesting that the interaction with CP190 is not critical for the *in vivo* functions of dCTCF.

## 2. Materials and Methods

### 2.1. Generation of CTCF^1^, CTCF^2^ point mutations and CTCF^attP^ deletions

CRISPR/Cas9-induced DNA double-strand breaks were used to generate *CTCF*^*1*^ and *CTCF*^*2*^ point mutations and a *CTCF* deletion. Guide RNAs (gRNAs) were selected using the program “CRISPR optimal target finder” (O’Connor-Giles Lab): C1 target: cttatggaaattgttgaggaagg; C2 target: aaaaaaggacgaggaccccg (for point mutations); 5’CTCFattP: aaaaaaggacgaggaccccg; 3’CTCF^attP^: gtgctggcttgattctctga (for deletion). To obtain *CTCF*^*1*^ and *CTCF*^*2*^ point mutations, CRISPR/Cas9 was combined with the non-homologous repair of double-stranded DNA breaks, followed by the selection of flies with a mutant dCTCF phenotype (Supplementary Figure 1). The F0 progeny was crossed with *y w*; *TM6/GE24185* flies, marked by the *mini-white* gene (yellow eyes). Selected males with a strong dCTCF mutant phenotype were crossed with *y w*; *TM6/MKRS* females, followed by the selection of flies without the *mini-white* reporter.

The deletion was obtained using CRISPR/Cas9-induced homologous recombination. The plasmid for homologous recombination were designed using a pBlu2SKP vector and contained several genetic elements, in the following order: [proximal arm]-[*lox*]-[*pAct5C-mCherry-SV40polyA*]-[*lox*]-[*attP*]-[distal arm] (Supplementary Figure 2). Homology arms were obtained by DNA amplification with the following primers: CTCF_5’f_d: gagggccccaagttctcggctgatca; CTCF_5’f_r: tcctcgagagcggatgaccgggat; CTCF_3’f_d: gagcggccgcagagaatcaagccagcac; CTCF_3’f_r: tcccgcgggtatgtacataagttagtcataa. The plasmid construct was injected into *58492* (Bloomington Drosophila Stock Center) embryos together with two gRNAs. The F0 progeny was crossed with *y w*; *TM6/MKRS* flies. Flies with potential deletions were selected based on *mCherry* fluorescence, and these flies were crossed with *y w*; *TM6/MKRS* flies. All independently obtained flies with the *mCherry* reporter were tested by PCR, and successful deletion events were confirmed by the sequencing of PCR products.

### 2.2. Fly crosses and transgenic lines

*Drosophila* strains were grown at 25°C under standard culture conditions. The transgenic constructs were injected into preblastoderm embryos using the φC31-mediated site-specific integration system [55]. The emerging adults were crossed with the *y ac w*^*1118*^ flies, and the progeny carrying the transgene were identified by *y*^*+*^ pigmented cuticle. Details of the crosses and primers used for genetic analysis are available upon request.

### 2.3. Cuticle preparations

Adult abdominal cuticles of homozygous enclosed 3-day old flies were prepared essentially as described in [56]. Photographs in the bright or dark field were taken on the Nikon SMZ18 stereomicroscope using Nikon DS-Ri2 digital camera, processed with ImageJ 1.50c4 and Fiji bundle 2.0.0-rc-46.

### 2.4. Plasmid construction

For *in vitro* experiments, protein fragments were either PCR-amplified using corresponding primers, or digested from dCTCF or CP190 cDNA and subcloned into pGEX-4T1 (GE Healthcare) or into a vector derived from pACYC and pET28a(+) (Novagen) bearing p15A replication origin, Kanamycin resistance gene, and pET28a(+) MCS.

To express 3xHA-tagged dCTCF and CP190 in the S2 cells, protein-coding sequences were subcloned into the pAc5.1 plasmid (Life Technologies). Details of the cloning procedures, primers, and plasmids used for plasmid construction are available upon request.

### 2.5. Pulldown assay

GST-pulldown was performed with Immobilized Glutathione Agarose (Pierce) in buffer C (20 mM Tris-HCl, pH 7.5; 150 mM NaCl, 10mM MgCl_2_, 0.1 mM ZnCl_2_, 0.1% NP40, 10% (w/w) Glycerol). BL21 cells co-transformed with plasmids expressing GST-fused derivatives of dCTCF and 6xHis-Thioredoxin-fused CP190 [1-126] and [245-606] were grown in LB media to an A600 of 1.0 at 37°C and then induced with 1 mM IPTG at 18°C overnight. ZnCl_2_ was added to final concentration 100 μM before induction. Cells were disrupted by sonication in 1ml of buffer C, after centrifugation lysate was applied to pre-equilibrated resin for 10 min at +4°C; after that, resin was washed four times with 1 ml of buffer C containing 500 mM NaCl, and bound proteins were eluted with 50 mM reduced glutathione, 100 mM Tris, pH 8.0, 100 mM NaCl for 15 min. 6xHis-pulldown was performed similarly with Zn-IDA resin (Cube Biotech) in buffer A (30 mM HEPES-KOH pH 7.5, 400 mM NaCl, 5 mM β-mercaptoethanol, 5% glycerol, 0.1% NP40, 10 mM Imidazole) containing 1 mM PMSF and Calbiochem Complete Protease Inhibitor Cocktail VII (5 μL/mL), washed with buffer A containing 30 mM imidazole, and proteins were eluted with buffer B containing 250 mM imidazole (20 min at +4°C).

### 2.6. Co-immunoprecipitation assay

*Drosophila* S2 cells were grown in SFX medium (HyClone) at 25°C. S2 cells grown in SFX medium were co-transfected by 3xHA-dCTCF (wild-type and with deletion of CP190-interacting region) and CP190 plasmids with Cellfectin II (Life Technologies), as recommended by the manufacturer. Protein extraction and co-immunoprecipitation procedure were performed as described in [57]. Anti-CP190 antibodies and rat IgG were used for co-immunoprecipitations. The results were analysed by Western blotting. Proteins were detected using the ECL Plus Western Blotting substrate (Pierce) with anti-HA and anti-CP190 antibodies.

### 2.7. Fly extract preparation

20 adult flies were homogenized with a pestle in 200 μL of 1xPBS containing 1% β-mercaptoethanol, 10 mM PMSF, and 1:100 Calbiochem Complete Protease Inhibitor Cocktail VII. Suspension was sonicated 3 times for 5 s at 5 W. Then, 200 μL of 4xSDS-PAGE sample buffer was added and mixture was incubated for 10 min at 100°C and centrifuged at 16,000 *g* for 10 min.

### 2.8. ChIP-Seq

Chromatin preparation from 2-3 day adult flies was performed as previously described [39, 40, 58]. Chromatin was precipitated with mouse anti-HA (1:100), rat anti-dCTCF (1;200), anti-CP190 (1:200) antibodies, or with nonspecific mouse and rat IgG. The ChIP-seq libraries were prepared with NEBNext® Ultra™ II DNA Library Prep kit, as described in the manufacturer’s instructions. Amplified libraries were quantified using fluorometry with DS-11 (DeNovix, United States) and Bioanalyzer 2100 (Agilent, United States). Diluted libraries were clustered on a pair-read flowcell and sequenced using a NovaSeq 6000 system (Illumina, United States). Raw and processed data were deposited in the NCBI Gene Expression Omnibus (GEO) under accession number GSE175402 (https://www.ncbi.nlm.nih.gov/geo/query/acc.cgi?acc=GSE175402; token for reviewer: utwngcqqfhcnfwp).

Chromatin immunoprecipitation sequencing (ChIP-seq) analysis was performed for six samples (dCTCF, HA, and CP190 in both the *dCTCF*^*wt*^ and *dCTCF*^*Δ705–733*^ lines); two biological replicates were obtained for each sample. Paired-end sequencing technology was applied, and the average read length was 101 bp. Reads were processed as described previously [59]. Briefly, the reads were trimmed with cutadapt and aligned to dm6 of the *Drosophila melanogaster* genome using Bowtie2 [60]. Duplicates and reads that mapped to blacklist regions were removed. Then, the reads were processed using the IDR pipeline (https://sites.google.com/site/anshulkundaje/projects/idr), where peak calling was performed for each replicate. The replicates and pseudoreplicates were merged using MACS version 2 against preimmune controls in paired-end mode (option format = BAMPE) [61]. A soft p-value threshold for MACS2 of 0.01 was used in IDR. The IDR threshold was set to 0.05 for true replicates and 0.01 for pseudoreplicates, according to the developers’ recommendations (https://hbctraining.github.io/Intro-to-ChIPseq/lessons/07_handling-replicates-idr.html). All samples showed good reproducibility (both the rescue ratio and the self-consistency ratio were less than 2). An optimal set of reproduced peaks was chosen for each sample for further analysis. Chip-seq coverage tracks (BedGraph) were obtained using deepTools [62] bamCoverage function, with a bin width of 100 bp and normalized using RPKM (Reads Per Kilobase of transcript, per Million mapped reads). Additionally, bamCoverage tracks were obtained using log fold change (LogFC) calculations compared against preimmune controls. *De novo* motif discovery was performed using ChIPMunk [63, 64]. For motif discovery, the top 500 peaks per sample were narrowed to those ±200 bp around the summit, and ChIPMunk was run in peak summit mode. Genome-wide motif sites were identified using sarus (https://github.com/VorontsovIE/sarus), with a p-value threshold of 1 × 10^−4^.

Further analysis was performed in R version 3.6.3 [65]. The colocalization and distribution of peaks between genome elements were analyzed using ChIPpeakAnno package version 3.20.1 [66] and Chipseeker version 1.26.0 [67]. After comparing the dCTCF peak sets obtained using anti-HA and anti-dCTCF-C antibodies (Supplementary Figure 5), we merged them and obtained a combined peak set for dCTCF in both the *dCTCF*^*wt*^ and *dCTCF*^*Δ705–733*^ lines. To investigate changes in the CP190 and dCTCF signals when the mutant Δ705–733 form of dCTCF was expressed, we performed linear regressions for CP190:

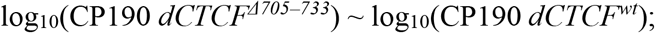

and dCTCF:

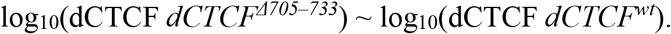

Then, we detected outliers by calculating studentized regression residuals, fitting a normal distribution to the residuals, and detecting peaks with a <5% probability of arising from the normal distribution (p-value < 0.05).

The dCTCF peaks that intersected with CP190 and those that did not were colocalized against other proteins known to interact with CP190 based on previously published data. For each of the proteins, we obtained raw reads and corresponding controls and processed them as described above, with minor deviations:

- If data was represented as single-end reads, Bowtie version 1 (Langmead et al. 2009) was used for mapping, and 200 bps was used as the MACS2 fragment width.
- If no replicates were available, the MACS2 p-value threshold was set to 0.00001.

The following proteins were used: Pita, Zinc finger protein interacting with CP190 (ZIPIC) [57], Su(Hw) [68], Ibf1, Ibf2 [44], and Insv [69]. For all proteins, only peaks with a corresponding motif site (sarus p-value < 0.0001) were used in the analysis.

## 3. Results

### 3.1. The inactivation of the dCTCF gene reduces the viability of adult flies and results in male and female sterility

Previous studies have disagreed on the viability of null *CTCF* mutants. Large deletions in the *dCTCF* gene are associated with lethality during the late pupae stage [30-32, 37]. However, the almost complete inactivation of *dCTCF* in the *GE24185* allele does not significantly affect the viability of *GE24185/GE24185* flies obtained from heterozygous parents [30, 36]. To explain these contradictory results, the *GE24185* allele was suggested to weakly express *dCTCF* at sufficient levels to enable the viability of adult flies. Alternatively, the existence of secondary mutations was hypothesized, which might enhance the expression of null mutations in the *dCTCF* gene.

To test these possibilities, we generated null *CTCF* alleles using a CRISPR/Cas9-based genome editing technology [53, 54]. We designed two independent experiments using different gRNA-recognized sequences in the N-terminal *dCTCF-*coding region (Fig. 1B, Supplementary Fig. 1).

We successfully obtained two independent mutant alleles (*CTCF*^*C1*^ and *CTCF*^*C2*^) that each features a single-base deletion resulting in a premature stop codon in the coding sequence of the *dCTCF* gene (Supplementary Fig. 1). Western blot analysis from adult males confirmed the absence of the dCTCF protein in *CTCF*^*C1*^ and *CTCF*^*C2*^ adult flies. Both mutants were complemented [resulting in a phenotype and viability similar to in wild-type (*wt*) flies] by one copy of the *Ubi:dCTCF* transgene (51D), which expresses cDNA encoding *dCTCF* under the control of the *Ubiquitin p63E* (*Ubi*) promoter (data not shown).

The previously characterized *dCTCF* mutations [30-33] predominantly affected the proper expression of *Abd-B*. The regulatory domains *iab-5, iab-6, iab-7*, and *iab-8* (Fig. 1A) control the gradual increase in the expression of *Abd-B* in segments A5, A6, A7, and A8, respectively [70, 71]. The boundaries (named *Mcp, Fab-6, Fab-7*, and *Fab-8*; Fig. 1A) provide functional autonomy to the *iab* domains during the regulation of *Abd-B* expression in the corresponding segments. Binding sites for dCTCF (Fig. 1A) have been identified in the *Mcp, Fab-6*, and *Fab-8* boundaries and in the *Abd-B* promoter region [30, 36, 72-75]. dCTCF is essential for the activity of the *Mcp, Fab-6*, and *Fab-8* boundaries [72, 76, 77].

Homozygous *CTCF*^*C1*^ and *CTCF*^*C2*^ mutants display phenotypes similar to that of *GE24185* flies (Fig. 2): held out wings, the presence of bristles on the male A6 sternite, and the deformation of genitalia. *GE24185* homozygous flies are characterized by delayed development and decreased viability, at 50% relative to *wt*. Homozygous *CTCF*^*C1*^ and *CTCF*^*C2*^ flies displayed low viability, around 10%. However, *CTCF*^*C1*^/*CTCF*^*C2*^ trans-heterozygotes showed viability similar to that of *GE24185* homozygotes. Thus, different secondary mutations can affect the viability of *CTCF*^*C1*^ or *dCTCF*^*C2*^ flies. Males from the *GE24185* and *CTCF*^*C2*^ lines show the partial transformation of A7 and A6 toward A6 and A5, respectively (Fig. 2). Rarely, males feature pigmented spots on the A4 tergite, suggesting that the *Mcp* boundary does not completely block crosstalk between the *iab-4* and *iab-5* enhancers, resulting in ectopic *Abd-B* expression in the A4 segment (data not shown). In contrast, *CTCF*^*C1*^ males present a relatively strong transformation of the A4 toward A5 (Fig. 2), suggesting that the *Mcp* functions are strongly affected. In *GE24185* females, the vaginal plates (A8) resemble those in *wt* flies; however, they are smaller in size. *GE24185* males and females are partially fertile and can produce progeny in crosses with *dCTCF*^*+*^ flies (Fig. 2). In contrast, *CTCF*^*C1*^ and *CTCF*^*C2*^ females and males are sterile and have stronger genital defects (Fig. 2). *GE24185/CTCF*^*C2*^ males and females and *CTCF*^*C1*^/*GE24185* males are partially fertile, whereas *CTCF*^*C1*^/*GE24185* females and both sexes of the *CTCF*^*C1*^*/CTCF*^*C2*^ line are sterile. Thus, the *CTCF*^*C1*^ chromosome contains an additional dominant mutation that affects fertility in the null *dCTCF* background.

**Figure 2.**
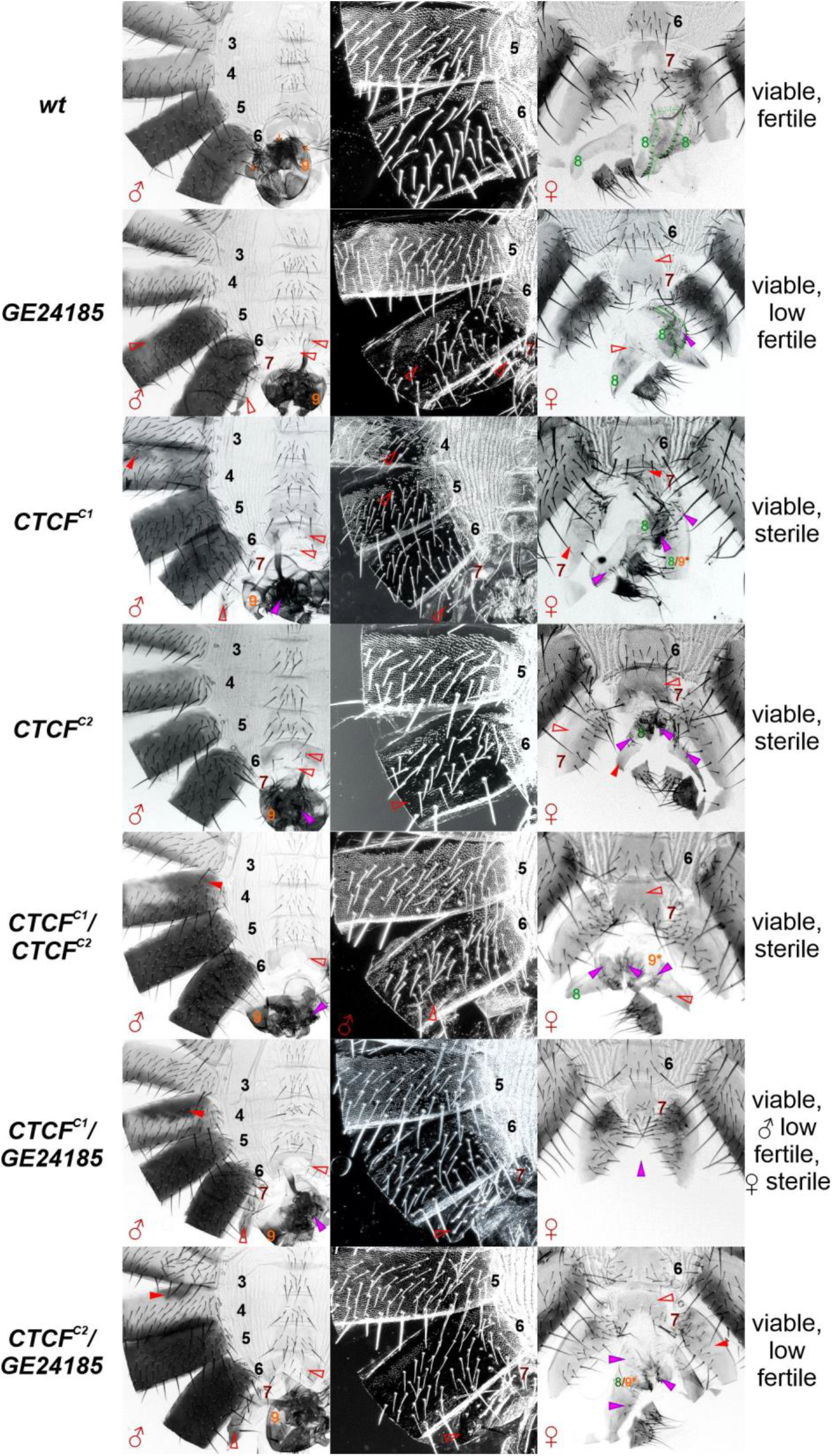
Morphology of the abdominal segments in the *dCTCF* mutations. The cuticles of the *wild-type* (*wt*) and mutant males are shown in the left and central (dark field) columns. The cuticles of the corresponding females are shown in the right column. The filled red arrowheads indicate morphological features indicative of gain of function (GOF) transformations associated with increased or ectopic *Abd-B* expression. The empty red arrowheads indicate signs of loss of function (LOF) transformations, associated with decreased *Abd-B* expression relative to that in *wt*. The purple arrowheads indicate signs of developmental disorders in the male and female genitalia. In *wt* males, *Abd-B* expression in the A5 and A6 segments is responsible for tergite pigmentation. The A5 sternite has a quadrilateral shape and is covered in bristles, and the tergite is covered with trichomes. The A6 segment displays a banana-shaped sternite without bristles, and no trichomes exist on the tergite except at the anterior and ventral edges. Segments A7 and A8 are completely missing. The A9 segment gives rise to the male genitalia (orange digit). In *wt* females, the A6 sternite has a quadrilateral form, covered with bristles. The size of the tergite in A7 decreases toward the center, and in section, it looks like a triangle, whereas the sternite in A7 is horseshoe-shaped, with bristles pointing toward the posterior. The A8 segment gives rise to tergites (thin depigmented plate without bristles) and vaginal plates (marked in green). The A9 segment is absent in adult females. In the *dCTCF* mutants, *Abd-B* expression decreased, which led to a predominance of LOF phenotypes. In males, the A6 sternite features bristles and tergites with several additional trichomes (LOF). Simultaneously, the partial inactivation of the *Mcp* boundary results in the ectopic activation of *Abd-B* in the A4 segment (GOF). In females, the A7 sternites became wider and acquired signs of an A6 phenotype (LOF). The expression of *Abd-B* in the A8–A9 segments was strongly affected in the *dCTCF* mutants: the male genitals were deformed, undeveloped, and sometimes duplicated; the female vaginal plates were reduced and undeveloped, and male-like genitalia structures were occasionally observed.

Next, we tested whether the deletion of the *dCTCF* gene has the same phenotypic effect as either the *CTCF*^*C1*^ or *CTCF*^*C2*^ point mutations. To address this question, we used a two-step genome engineering platform that combines CRISPR-mediated homology-directed repair with φC31 [78]. A 2,173-bp region of the *dCTCF* gene, from the first intron to the end of third exon (3L:7,352,322…7,358,209, FlyBase, R6) was substituted with the *Act5C:mCherry* reporter, flanked by *lox* sites and containing an *attP* site (*CTCF*^*attP(mCh)*^, Fig. 1B, Supplementary Fig. 2).

Flies homozygous for the *CTCF*^*attP(mCh)*^ allele died at the late pupae stage and as pharate adults. After the deletion of the *Act5C:mCherry* reporter by the induction of recombination between the *lox* sites, the resulting *CTCF*^*attP*^ flies died at the same stages as the *CTCF*^*attP(mCh)*^ flies. The *Ubi:dCTCF* (51D) transgene was able to restore viability but not fertility, resulting in a nearly *wt-*like phenotype in the *CTCF*^*attP(mCh)*^ flies, although a pigment spot and several bristles on the A6 sternite were detected (Supplementary Fig. 3). The survival of *CTCF*^*attP*^ homozygotes with a *Ubi:dCTCF* background was much worse and decreased in the subsequent generations. The re-integration of the *dCTCF* gene in the *CTCF*^*attP*^ line completely restored both viability and fertility. The deletion of the *dCTCF* gene in *CTCF*^*attP*^ flies appeared to affect the expression of the surrounding genes, enhancing the mutant phenotype.

*CTCF*^*attP*^ males displayed strong ectopic pigmentation of the A4 segment (A4→A5 transformation), similar to that observed in *CTCF*^*C1*^ males (Supplementary Fig. 4). Similar ectopic activation of *Abd-B* was observed in *CTCF*^*attP*^*/CTCF*^*C1*^ males but not in *CTCF*^*attP*^*/GE24185* or *CTCF*^*attP*^*/CTCF*^*C2*^ males, suggesting that additional mutations affected the *Mcp* boundary functions in the *dCTCF*^*−*^ background. Taken together, these results suggested that many different alleles can affect the expressivity of the null *dCTCF* mutations.

### 3.2. Mapping of the dCTCF region that interacts with CP190

CP190 is known to be the primary partner of the dCTCF protein [45]. CP190 (Fig. 3A) has an N-terminal BTB domain, followed by an aspartate-rich D domain, a microtubule-targeting (M) domain, four C2H2 zinc fingers, and a glutamate-rich C-terminal domain [79]. Previously, we demonstrated through yeast two-hybrid (Y2H) and co-immunoprecipitation (co-IP) studies that the BTB and M domains of CP190 interact with an approximately 200-aa region in the C-terminal domain of dCTCF [36]. To map the interacting regions in the proteins more precisely, we used a glutathione-S-transferase (GST) pulldown assay. We tagged different regions of the C-terminal dCTCF domain, the CP190 BTB domain (Fig. 3B), and the CP190 M domain (Fig. 3C) with either GST or thioredoxin-6×His-tag. The pulldown experiments showed that the 715– 733 aa region of dCTCF was necessary for the interaction with the BTB domain of CP190.

**Figure 3.**
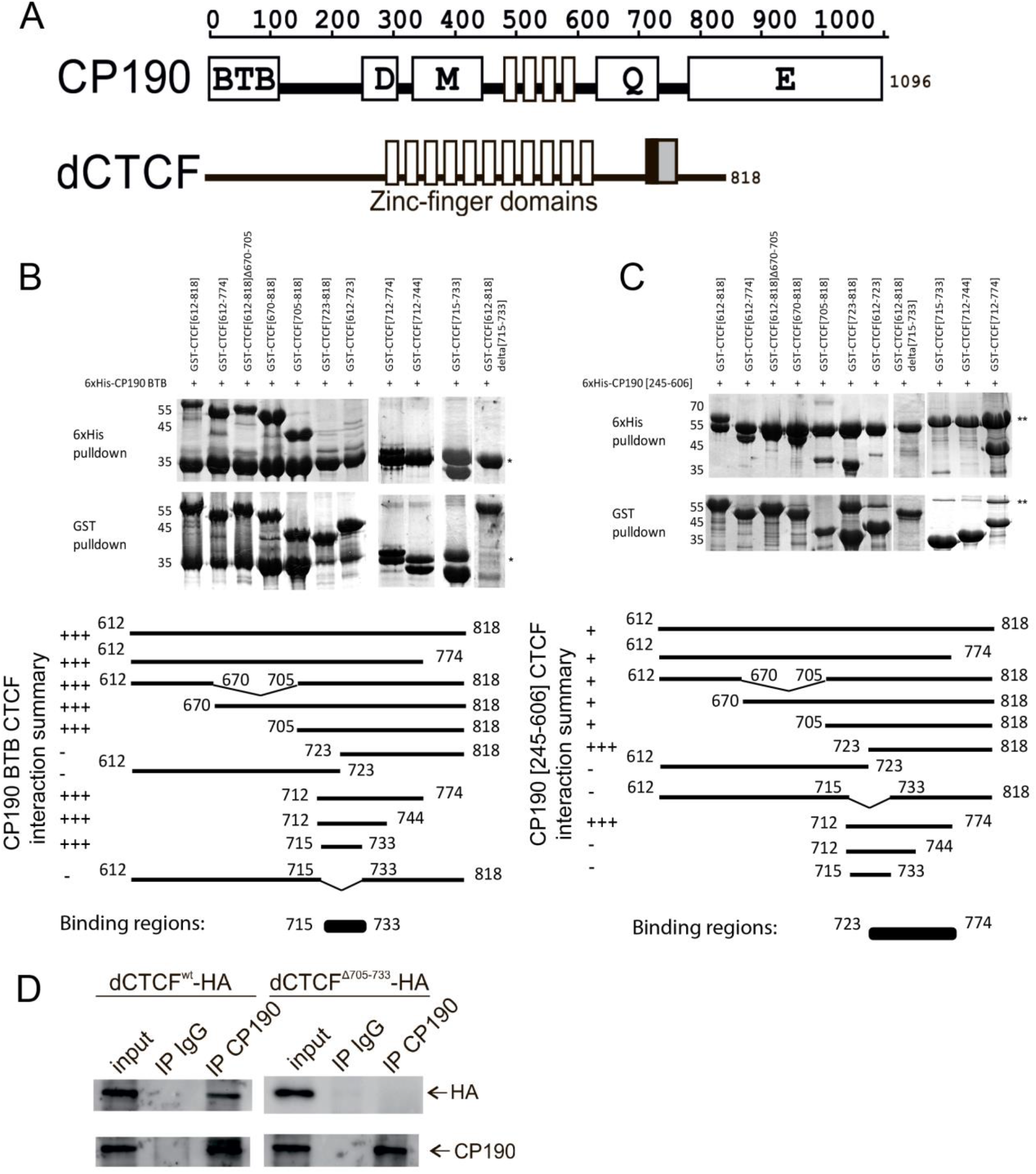
Mapping the CP190-interacting regions in the dCTCF protein. (A) A schematic representation of the full-length CP190 and dCTCF proteins. The interacting regions in the proteins are indicated by black (involving the BTB domain) and gray (involving the M domain) boxes. The positions of the amino acid residues in the proteins are shown above the diagrams. (B) GST and 6×His pulldowns of GST-fused dCTCF protein fragments co-expressed with the thioredoxin-6×His-fused CP190 BTB domain. The positions of the amino acids are given in square brackets. (C) GST and 6×His pulldowns of GST-fused dCTCF protein fragments co-expressed with the thioredoxin-6×His-fused CP190 M domain [245–606]. A schematic summary of the pulldown results. (D) Co-immunoprecipitation of CP190 with dCTCF^wt^ or dCTCF^Δ705–733^ protein fused with 3×HA in S2 cells. Protein extracts from *Drosophila* S2 cells cotransfected with 3×HA-dCTCF and CP190 plasmids were immunoprecipitated with antibodies against CP190 or nonspecific IgG (as a negative control), and the immunoprecipitates (IP) were analyzed by western blotting for the presence of 3×HA-tagged dCTCF proteins. The quality of the IP was controlled by western blotting for the presence of CP190 protein. “Input” refers to samples of the initial protein extract; “output” refers to the supernatant after the removal of the IP.

Interestingly, the M domain of CP190 interacted with a nearby sequence of dCTCF, which was mapped between 723 and 774 aa. The deletion of the 715–733 aa region from the C-terminal domain prevented the interaction with both CP190 domains.

To confirm these results *in vivo*, we expressed 3×HA-tagged dCTCF^wt^ and dCTCF^Δ705–733^ in S2 cells under the control of the ubiquitin promoter (Fig. 3D). We found that dCTCF but not dCTCF^Δ705–733^ interacted with the CP190 protein in co-IP experiments. These results confirmed the importance of the 715–733 aa region for the interaction between dCTCF and CP190.

### 3.3. Interaction with CP190 is not critical for the in vivo functions of dCTCF protein

The interaction between CP190 and dCTCF has been suggested to be necessary for the organization of distance interactions [80, 81]; therefore, the consequences of deleting the dCTCF domain necessary for CP190 interaction were explored. We used the *attP* site in the *CTCF*^*attP(mCh)*^ allele to insert the gene encoding *wt* (*dCTCF*^*wt*^*-HA*) or mutant (*dCTCF*^*Δ705–733*^*-HA*) dCTCF, with an HA tag at the C-terminus. The insertion included all *CTCF* genome regions from the first intron to the fourth exon, with the HA tag incorporated within the fourth exon, and a polyadenylation signal from SV40 and the *white* reporter flanked by *lox* sites (Supplementary Fig. 2). After the deletion of the reporter genes, the only *lox* site is present in the first intron of the d*CTCF* gene. d*CTCF*^*Δ705–733*^ flies displayed a *wt* phenotype and normal fertility. Western blot analysis showed that dCTCF^wt^ and dCTCF^Δ705–733^ are expressed at similar levels, comparable to the expression level of dCTCF in *Oregon* line (Fig. 4). These results suggested that the direct interaction with CP190 is not critical for the functional activity of dCTCF *in vivo*.

**Figure 4.**
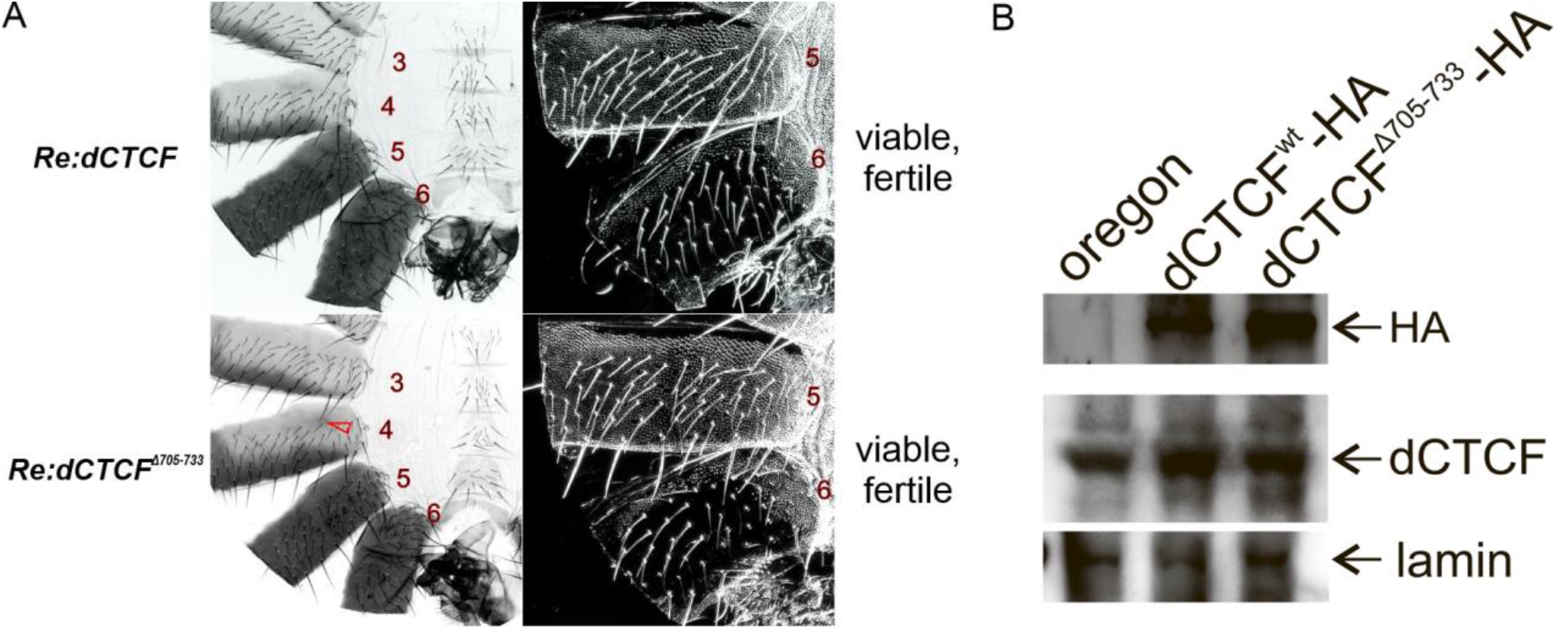
Mutations in the endogenous *dCTCF* gene. (A) Morphologies of the abdominal segments in *CTCF*^*wt*^ and *CTCF*^*Δ705–733*^ males. All other designations are the same as in Fig. 2. (B) Immunoblot blot analysis (10% SDS PAGE) of protein extracts prepared from adult 2-day-old males of the *Oregon, CTCF*^*wt*^ and *CTCF*^*Δ705–733*^ lines with dCTCF and lamin (internal control) antibodies.

### 3.4. Interaction with CP190 is not required for dCTCF to bind to chromatin

The next task was to study the level of interdependence between dCTCF and CP190 binding and interactions with chromatin. For this goal, we compared the binding of the dCTCF and CP190 proteins in transgenic lines expressing either dCTCF^wt^-HA or dCTCF^Δ705–733^-HA. To map the chromatin-binding sites of CP190 and dCTCF-HA, we obtained chromatin collected from two-day-old flies.

For each transgenic line, we performed ChIP using previously characterized antibodies, followed by ChIP-seq using Illumina’s massive parallel sequencing technology. For dCTCF, we used previously characterized antibodies against the C-terminal domain (α-CTCF, 612–818) [36] and antibodies against the HA×3 epitope. For CP190, we used antibodies (α-CP190) against the region between 308–1096 aa [36]. The comparison of specific ChIP patterns resulted in the identification of 453 HA×3 peaks and 462 dCTCF peaks, with motifs in chromatin isolated from dCTCF^wt^-HA adults (Supplementary Fig. 5A and D). The CTCF-binding profiles obtained using the HA and α-CTCF antibodies were highly reproducible. In both experiments, we identified the same motif that completely matched a previously identified motif for dCTCF [26, 28]. In total, 422 peaks were identical in both experiments (approximately 95%); therefore, we summarized peaks using the motifs from both experiments. When we correlated the distribution of these peaks with genomic annotation, we found that they were concentrated in the promoter and intron regions. Similar results were obtained using a transgenic line expressing dCTCF^Δ705–733^-HA (Supplementary Fig. 5A), in which 82% of dCTCF sites overlapped between dCTCF^wt^-HA and dCTCF^Δ705–733^-HA. The peaks for dCTCF without motifs demonstrated poor reproducibility for both the HA and dCTCF antibodies and weak binding efficiency with chromatin (Supplementary Fig. S5B and C and Supplementary Fig. 6).

To investigate changes in chromatin binding for CP190 and dCTCF in the transgenic line expressing dCTCF^Δ705–733^, ChIP-seq signal values were estimated in the set of α-CTCF peaks, with motifs reproduced in dCTCF^wt^ and dCTCF^Δ705–733^ adults (Fig. 5A). From these 575 peaks, we identified 40 peaks that demonstrated an enhanced signal in dCTCF^wt^ adults compared with dCTCF^Δ705–733^ adults (Fig. 5C). As a result, the dCTCF^Δ705–733^ binding efficiency was only significantly reduced for a few genome sites (Fig. 5C and D). Thus, CP190 binding is not essential for dCTCF binding to most chromatin sites.

**Figure 5.**
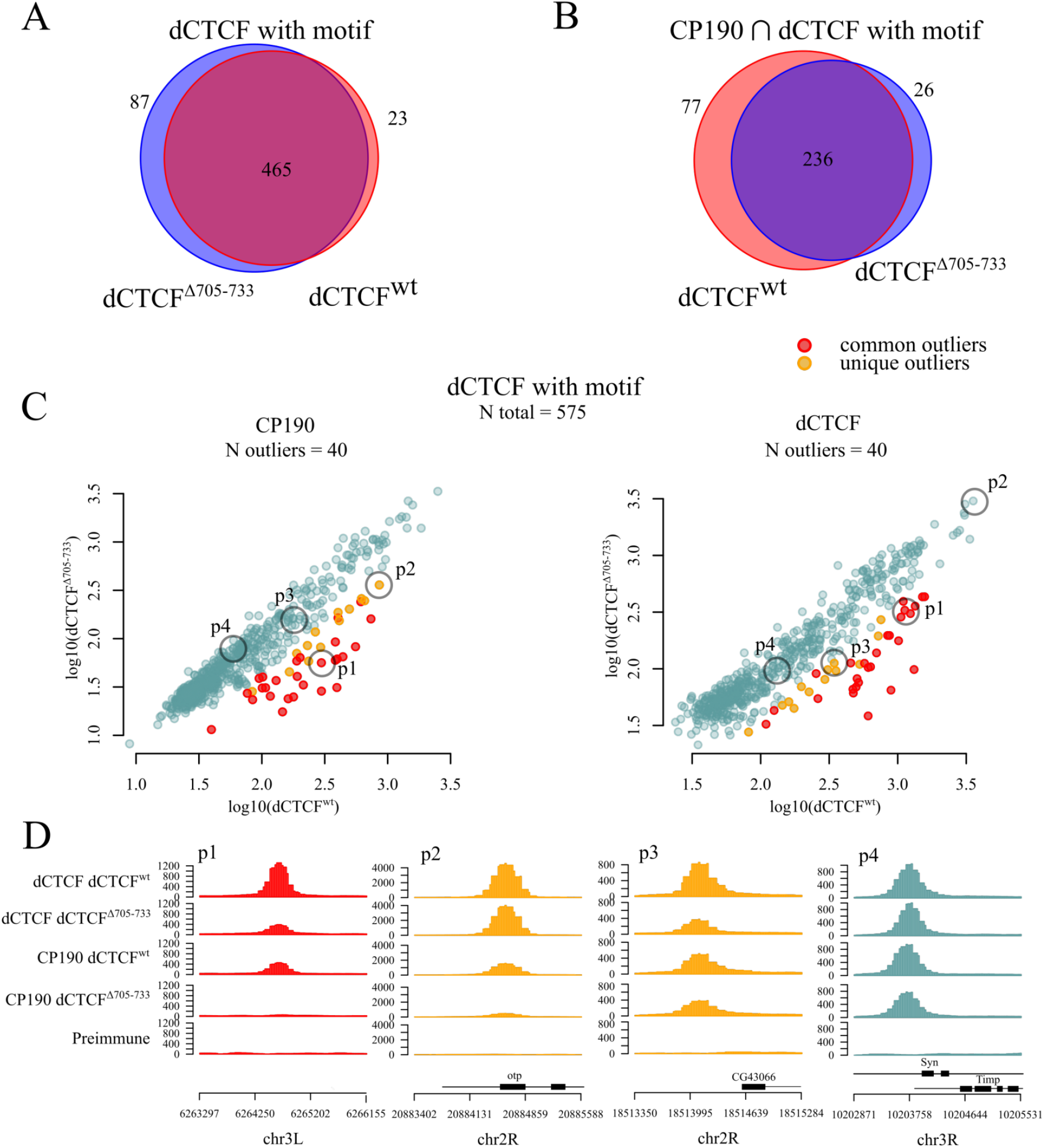
Differences in the dCTCF and CP190 ChIP-Seq results between the *dCTCF*^*Δ705–733*^ and *dCTCF*^*wt*^ lines. Only dCTCF peaks associated with dCTCF motif sites were considered. (A) Venn diagram showing the number of overlapping dCTCF peaks between the *dCTCF*^*Δ705–733*^ and *dCTCF*^*wt*^ lines. (B) Venn diagram showing the numbers of overlapping CP190 and dCTCF peaks between the *dCTCF*^*Δ705–733*^ and *dCTCF*^*wt*^ lines. (C) Comparison of the dCTCF peaks obtained from the dCTCF and CP190 ChIP-Seq signals (log10(RPKM)) in different lines (*dCTCF*^*Δ705–733*^ and *dCTCF*^*wt*^ merged). Outliers were determined using linear regression (see Methods). Peaks that were determined to be outliers in both the CP190 and dCTCF signal analyses were denoted as “common outliers” (red). Peaks determined as outliers only for one of the proteins were denoted as “unique outliers” (yellow). (D) Examples of protein signals (RPKM) in peaks that appeared to be common outliers (red), unique CP190 outliers (yellow), unique dCTCF outliers (yellow), and not an outlier (blue). In all visualizations, only those dCTCF peaks associated with dCTCF motif sites were used.

We compared the CP190 signal between the dCTCF^wt^/CP190 and dCTCF^Δ705–733^/CP190 groups of peaks and found that 313 of 488 dCTCF^WT^-HA peaks and 262 of 552 dCTCF^Δ705–733^-HA peaks overlapped with CP190 (Fig. 5B). The partial weakening of CP190 binding was observed at 40 sites, although almost all sites demonstrated the maintenance of stable CP190 binding (Fig.5 C and D). These results suggested that dCTCF is not essential for recruiting CP190 to chromatin. The independence of CP190 chromatin binding is likely due to dCTCF colocalizing with the binding sites of other proteins, which can also recruit CP190. Approximately 45% of dCTCF+CP190 binding sites in promoter regions colocalized with the binding sites for Su(Hw), Ibf1, Ibf2, Insv, Pita, or ZIPIC, which can all interact with CP190 (Supplementary Fig. 7) [27, 39, 44, 68, 82].

## 4. Discussion

Our current and previous results [30-32, 37, 83] showed that flies homozygous for independently obtained null *CTCF* mutations have different phenotypic manifestations and that all null *CTCF* mutations could be complemented by the *dCTCF*^*+*^ transgene. Thus, many different alleles of unknown genes associated with the third chromosome appear to affect *dCTCF* mutation expressivity. Previously, dCTCF was demonstrated as non-essential for mutants to progress through embryogenesis and larval stages [32]. Here, we demonstrated that null *dCTCF* mutants are viable into adulthood. The absence of any strong effects on the development of *Drosophila* in null *dCTCF* mutants is likely due to cooperation with other architectural proteins. The dCTCF protein is considered essential for the expression of the *Abd-B* gene, which correlates with the presence of CTCF-binding sites in almost all boundaries, with the exception of *Fab-7*. The inactivation of dCTCF is associated with decreased *Abd-B* expression in the A6, A7, A8, and A9 segments and the partial sterility of males and females. dCTCF is likely required for effective communication among the *iab-6, iab-7*, and *iab-8* enhancers and the *Abd-B* promoters. The ectopic activation of *Abd-B* in the A4 segment observed in *dCTCF* null mutants suggests that dCTCF is essential for the insulator function of the *Mcp* boundary.

Here, we precisely mapped the CP190 interacting regions in the C-terminal segment of dCTCF. The BTB and M domains of CP190 bind with the partially overlapping dCTCF regions 715–733 aa and 723–744 aa, respectively. The simultaneous interaction between dCTCF and two CP190 domains likely improves the co-binding of these proteins to chromatin. Previously, dCTCF binding to chromosomes was thought to be dependent on CP190 interaction [30, 37]. In contrast with this belief, our results demonstrated that the dCTCF^Δ705–733^ mutant, which lacks the CP190 interacting domain, was able to bind to chromosomes with the same efficiency as the wild-type protein.

CP190 is an essential chromatin-binding protein that mediates the binding of several transcriptional complexes involved in chromatin opening, including NURF, SAGA, and dREAM, and recruits the histone methyltransferase dMes4 to dCTCF-binding sites [47, 49, 52, 84].CP190 has also been speculated as being necessary for the organization of distance interactions between various chromatin sites bound to architectural proteins [41, 81, 85]. Moshkovich et al.[33] showed that dCTCF together with CP190 are responsible for the formation of chromatin loops between boundaries in BX-C. The mutant dCTCF protein lacking the CP190 interacting domain (CTCF^Δ705–733^) remained fully functional in the regulation of *Abd-B* expression. By contrast, the dCTCF mutant generated by the deletion of the N-terminal dimerization domain was unable to support *Abd-B* expression, similar to the null *dCTCF* mutations [36]. The presence of the N-terminal dimerization domain in the dCTCF^Δ705–733^ mutant may explain its ability to support normal *Abd-B* expression.

The boundaries in *BX-C* are formed by binding sites for several different architectural proteins, including dCTCF, Pita, and Su(Hw) [10]. Other chromatin-binding proteins appear to compensate for the inability of CTCF^Δ705–733^ to recruit the CP190 protein to the *BX-C* boundary and across the genome. Approximately 45% of the dCTCF-CP190 peaks colocalized with other known CP190-interacting proteins.

The comparison of recent CTCF ChIP and Hi-C data between *Drosophila* and humans suggested that dCTCF contains a relatively small number of binding sites in comparing with hCTCF and does not play a major role in the organization of the genome architecture [20, 50, 86, 87]. However, dCTCF is enriched at TAD borders in a cell line derived from the larval central nervous system [88]. Thus, dCTCF might be involved in the organization of TADs that are specific to particular *Drosophila* tissue types. Approximately half of all dCTCF sites colocalized with cohesin, which suggests the possibility of interplay between cohesion and dCTCF during the organization of chromatin loop domains [89]. In support of this possibility, dCTCF contains a motif similar to that found to be important for the interaction between hCTCF and cohesion [23]. Therefore, the primary difference between hCTCF and dCTCF appears to be the number of binding sites in the genome, resulting in different contribution levels to the organization of the chromosomal architecture in humans and flies. In all organisms, CTCF belongs to a large group of proteins that cooperatively organize chromatin architecture, and further research remains necessary to clarify the contributions made by all of these proteins. The absence of an essential role for dCTCF in the development of *Drosophila* will allow for the functional roles of individual domains in this protein to be studied, including the degree of their evolutionary conservatism.

**Supplementary data** to this article can be found online at

## Transparency document

The Transparency document associated with this article can be found, in online version.

## Acknowledgements

We thank Farhod Hasanov and Aleksander Parshikov for fly injections. We are grateful to the Center for Precision Genome Editing and Genetic Technologies for Biomedicine IGB RAS for bioinformatics.

## Author contribution statement

O.M., A.B., N.Z., N.P., O.K. performed the genetics experiments,

O.M. performed ChIP-Seq, CoIP and Western blot analysis

A.B. performed biochemistry.

N.K. carried out the bioinformatic work.

O.M. and O.K. evaluated and analyzed the data, prepare figures and wrote the manuscript.

P.G. planned the project, evaluated and analyzed the data, and wrote the manuscript.

## Supplementary Information

**Figure 1.**
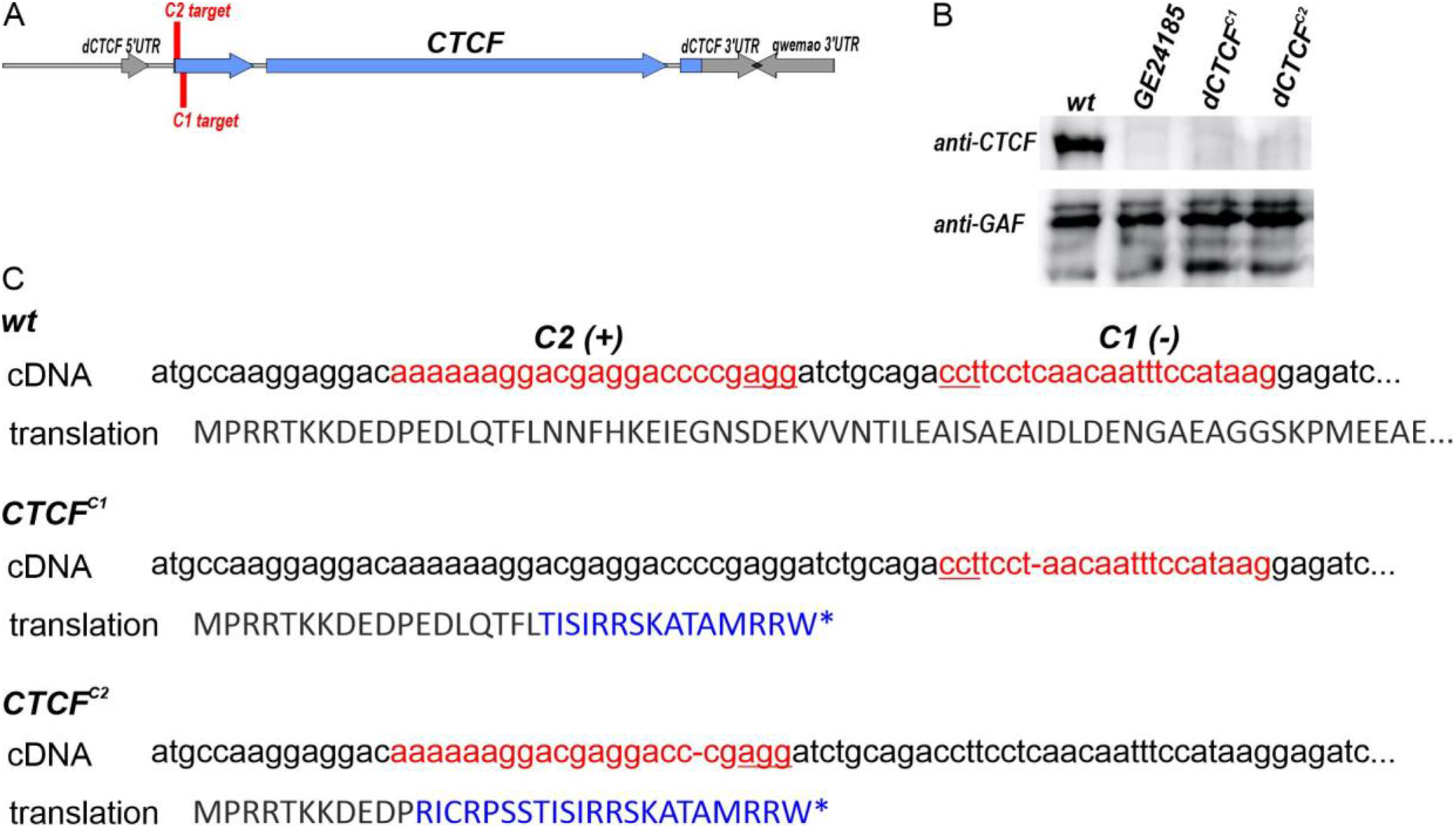
Generation of the null *dCTCF* mutations. (A) The map of the *dCTCF* genome region. The *dCTCF*-coding regions are shown as blue horizontal arrows. The surrounding genes (*pst* and *qwemao*) and untranslated regions of the *dCTCF* gene are shown as gray arrows. CRISPR targets are shown as vertical red bars (C1 and C2). (B) Western blotting analysis of the protein extracts prepared from adult 2-day-old males expressing the wild-type *dCTCF* gene (*wt)* or any of the *dCTCF* mutations: *GE24185, CTCF*^*C1*^, and *CTCF*^*C2*^. Immunoblot analysis was performed with anti-dCTCF and anti-GAGA factor (GAF, internal control) antibodies. (C) The nature of the *CTCF*^*C1*^and *CTCF*^*C2*^ mutations. The C1 and C2 guides are shown with red letters. The deleted nucleotides in the *CTCF*^*C1*^, *CTCF*^*C2*^ are shown as “−. “ Blue capital letters indicate the appearance of new amino acids after changes in the open reading frame (ORF). The last amino acids in the *dCTCF* mutations are marked with asterisks.

**Figure 2.**
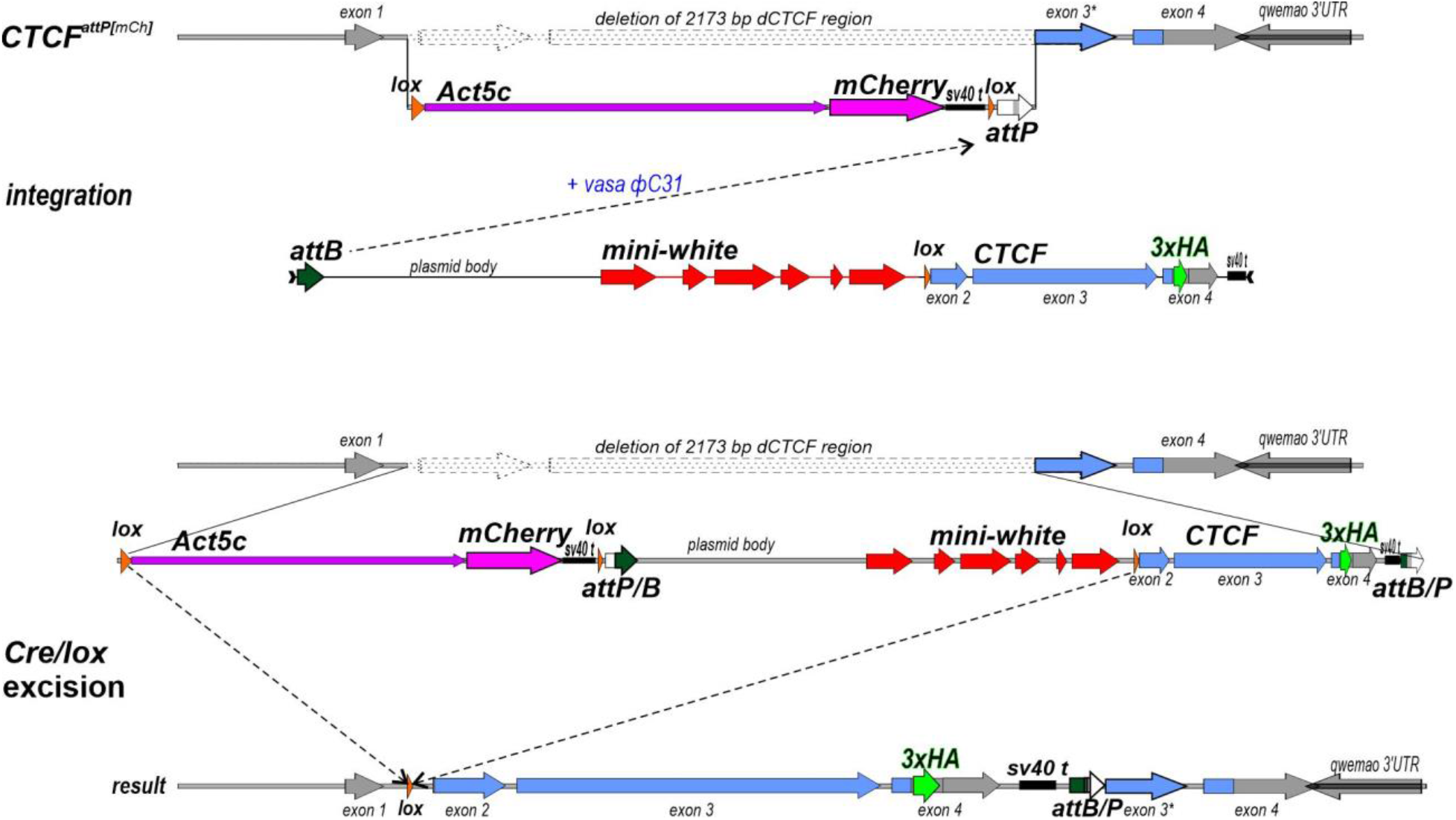
Strategy for generating the *CTCF*^*wt*^ and *CTCF*^*Δ705–733*^ alleles. In the *CTCF*^*attP(mCh)*^ platform, the 2,173-bp *CTCF* gene, from the first intron to the end of the third exon (3L:7,352,322…7,358,209, FlyBase, R6), was replaced with the *attP-Act5c-mCherry*. The *dCTCF*-coding regions and the untranslated regions (UTRs) are shown as blue and gray horizontal arrows, respectively. The surrounding genome regions are indicated as gray lines. The *mCherry* reporter (magenta arrow) is controlled by the *Act5C* promoter (green arrow). The *attP* and *lox* sites used for genome manipulations are shown as white and orange arrows, respectively. The construct for the replacement contains the mini-white reporter, a *lox* site, and the *CTCF*-coding region fused with a 3×HA tag. The attP-mediated recombination results in the integration of a complete construct, including the plasmid DNA, into the *CTCF*^*attP(mCh)*^ platform. Cre-mediated recombination between the *lox* sites generates the deletion of all sequences, except for the *dCTCF* gene and one *lox* site in the first intron.

**Figure 3.**
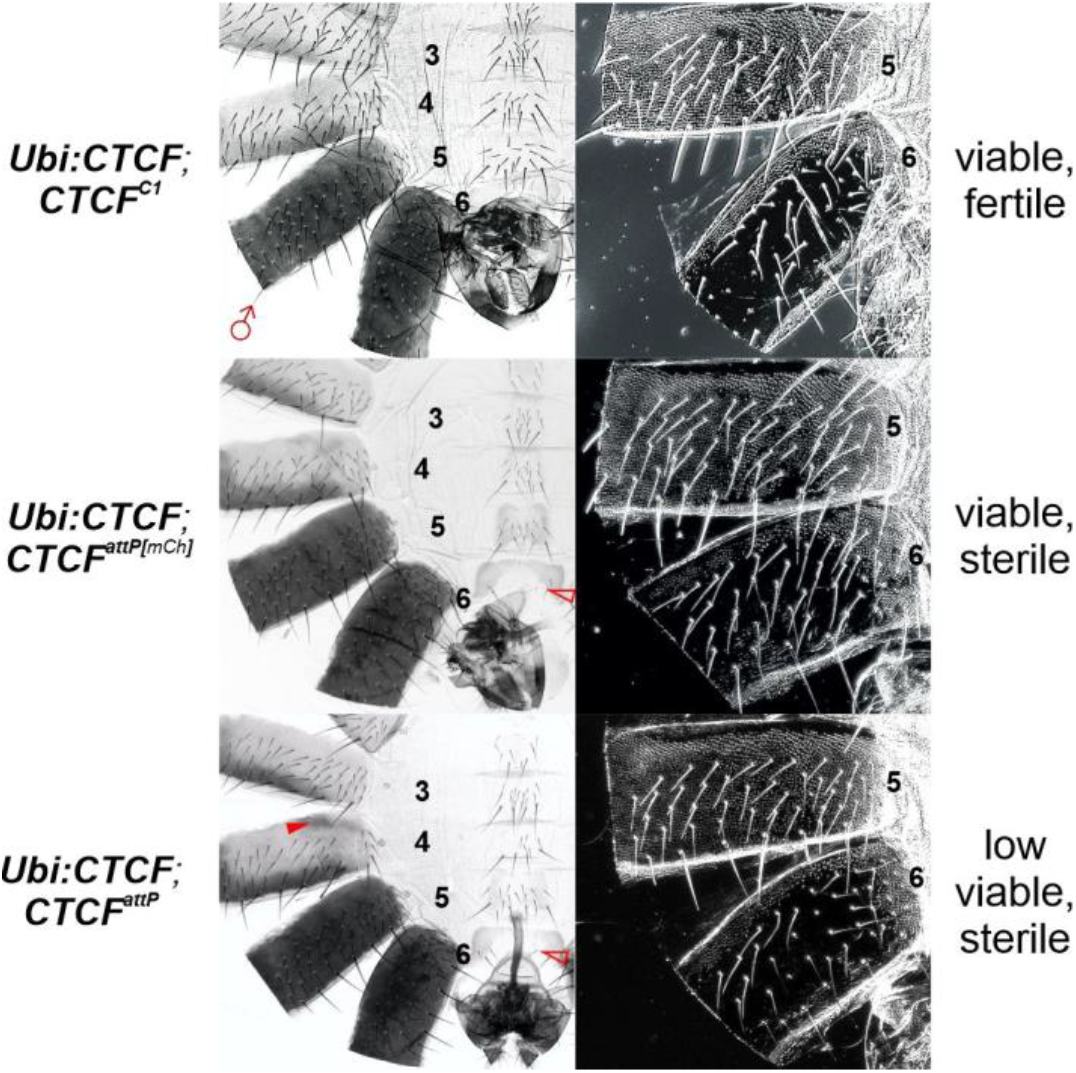
Morphology of the abdominal segments in the *CTCF* mutations, when combined with the *Ubi:CTCF* (51D) transgene. The filled red arrowheads show the morphological features indicative of gain of function (GOF) transformations associated with increased *Abd-B* expression relative to WT. The empty red arrowheads show the signs of loss of function (LOF) transformations associated with decreased *Abd-B* expression.

**Figure 4.**
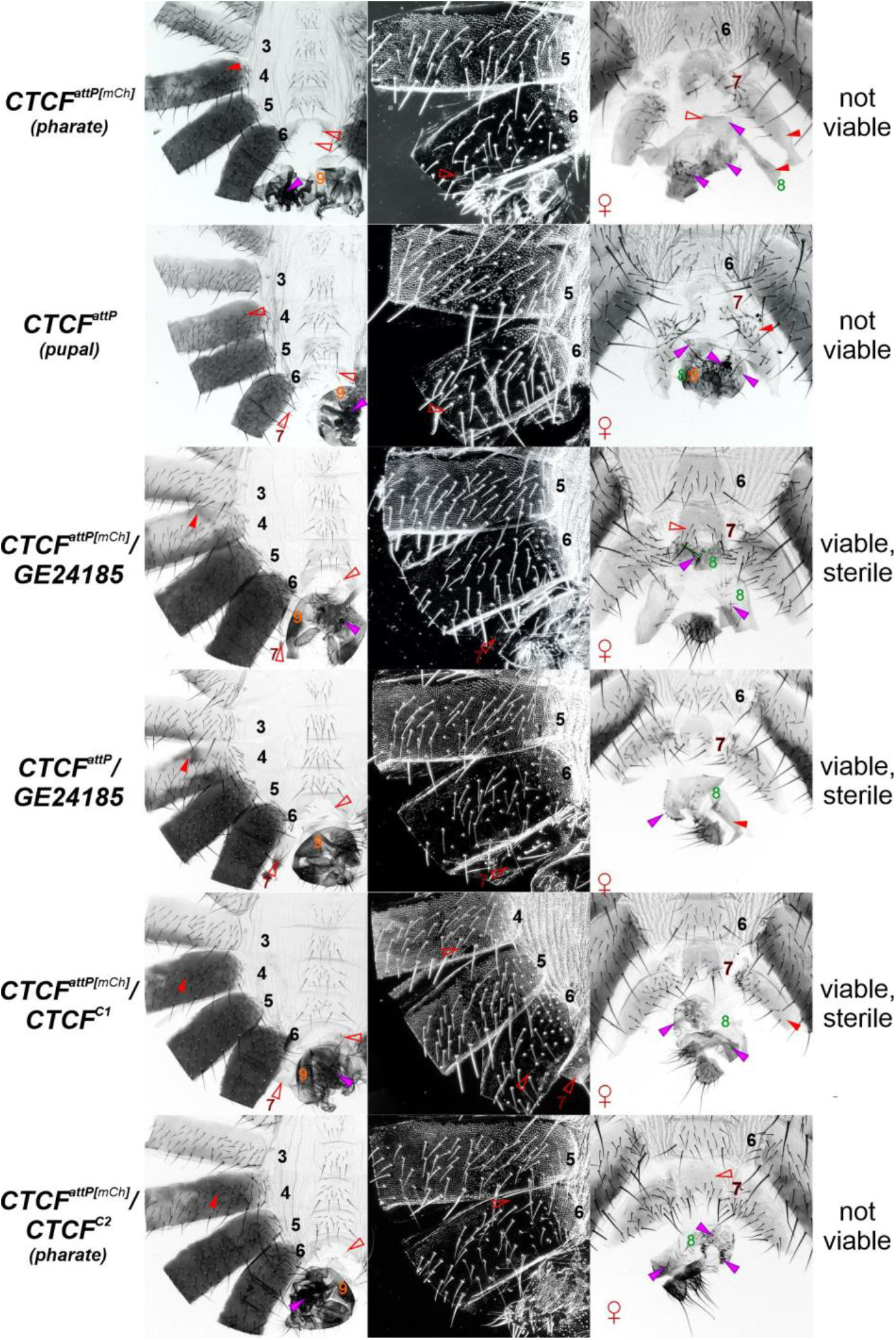
Morphology of abdominal segments in *CTCF*^*attP(mCh)*^ flies and derivatives. The cuticles of the *wild-type* (*wt*) and mutant males are shown in the left and central (dark field) columns. The cuticles of corresponding females are shown in the right column. The filled red arrowheads indicate morphological features that represent gain of function (GOF) transformations associated with increased or ectopic *Abd-B* expression. The empty red arrowheads indicate the signs of loss of function (LOF) transformations associated with decreased *Abd-B* expression relative to *wt*. The purple arrowheads indicate signs of developmental disorders in the male and female genitalia. In *wt* males, *Abd-B* expression in the A5 and A6 segments is responsible for tergite pigmentation. The A5 sternite has a quadrilateral shape and is covered in bristles, and the tergite is covered with trichomes. The A6 segment forms a banana shape in the sternite without bristles, and tergites do not feature trichomes except at the anterior and ventral edges. Segments A7 and A8 are completely missing. The A9 segment gives rise to male genitalia (orange digit). In *wt* females, the A6 sternite has a quadrilateral form and is covered with bristles. The size of tergites in A7 decreases toward the center, and in section, it resembles a triangle, whereas the sternite in A7 is horseshoe-shaped with bristles pointing toward the posterior. Segment A8 gives rise to tergites (thin depigmented plates without bristles) and vaginal plates (marked in green). The A9 is absent in adult females. In the *dCTCF* mutants, the expression of *Abd-B* is decreased, resulting in a mixture of LOF and GOF phenotypes. In males, the A6 sternite features bristles and tergites with several additional trichomes (LOF). Simultaneously, the partial inactivation of the *Mcp* boundary results in the ectopic activation of *Abd-B* in the A4 segment (GOF). In females, the A7 sternites become wider and acquired signs of an A6 phenotype (LOF). The expression of *Abd-B* in the A8–A9 segments is strongly affected in *<u>dCTCF</u>* mutants: the male genitals are deformed, undeveloped, and sometimes duplicated; the female vaginal plates are reduced and undeveloped, and male-like genitalia structures are sometimes observed.

**Figure 5.**
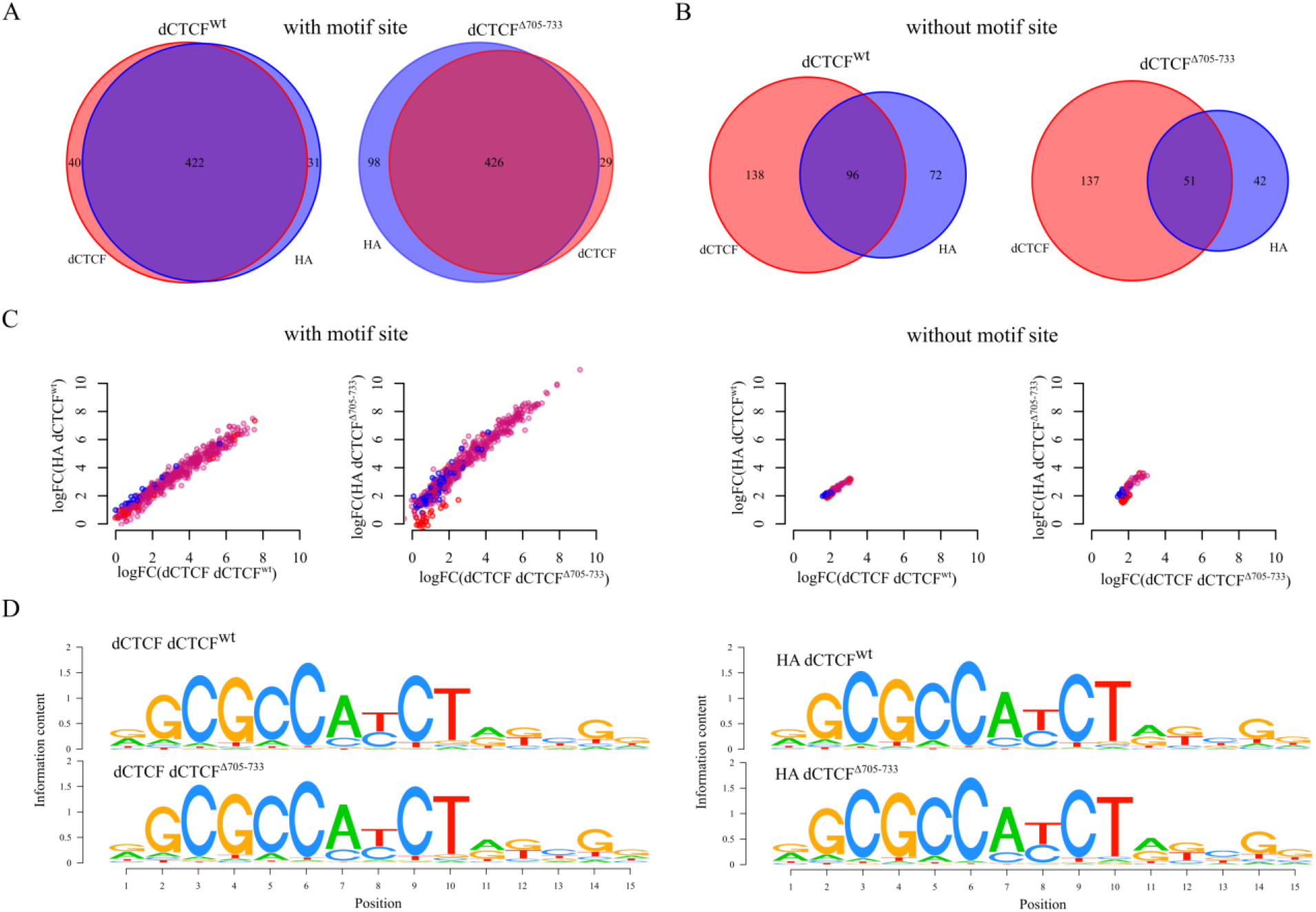
Comparison between the dCTCF-binding sites revealed using HA and dCTCF antibodies in *dCTCF*^*Δ705–733*^ and *dCTCF*^*wt*^ lines. (A and B) Venn diagrams showing overlaps between dCTCF peaks observed when using HA and dCTCF antibodies for those containing dCTCF motif sites (A) and those without motif sites (B). (C) Comparison of the dCTCF signals (LogFC between target signal and preimmune control signal) identified using the HA and dCTCF antibodies and dCTCF-binding sites (HA and dCTCF merged). Peaks that were only revealed using HA antibodies are colored in red, and peaks that were only revealed using dCTCF antibodies are in blue, and common peaks are colored in purple. (D) Sequence logos for dCTCF peaks revealed using HA and dCTCF antibodies in *dCTCF*^*Δ705–733*^ and *dCTCF*^*wt*^ lines.

**Figure 6.**
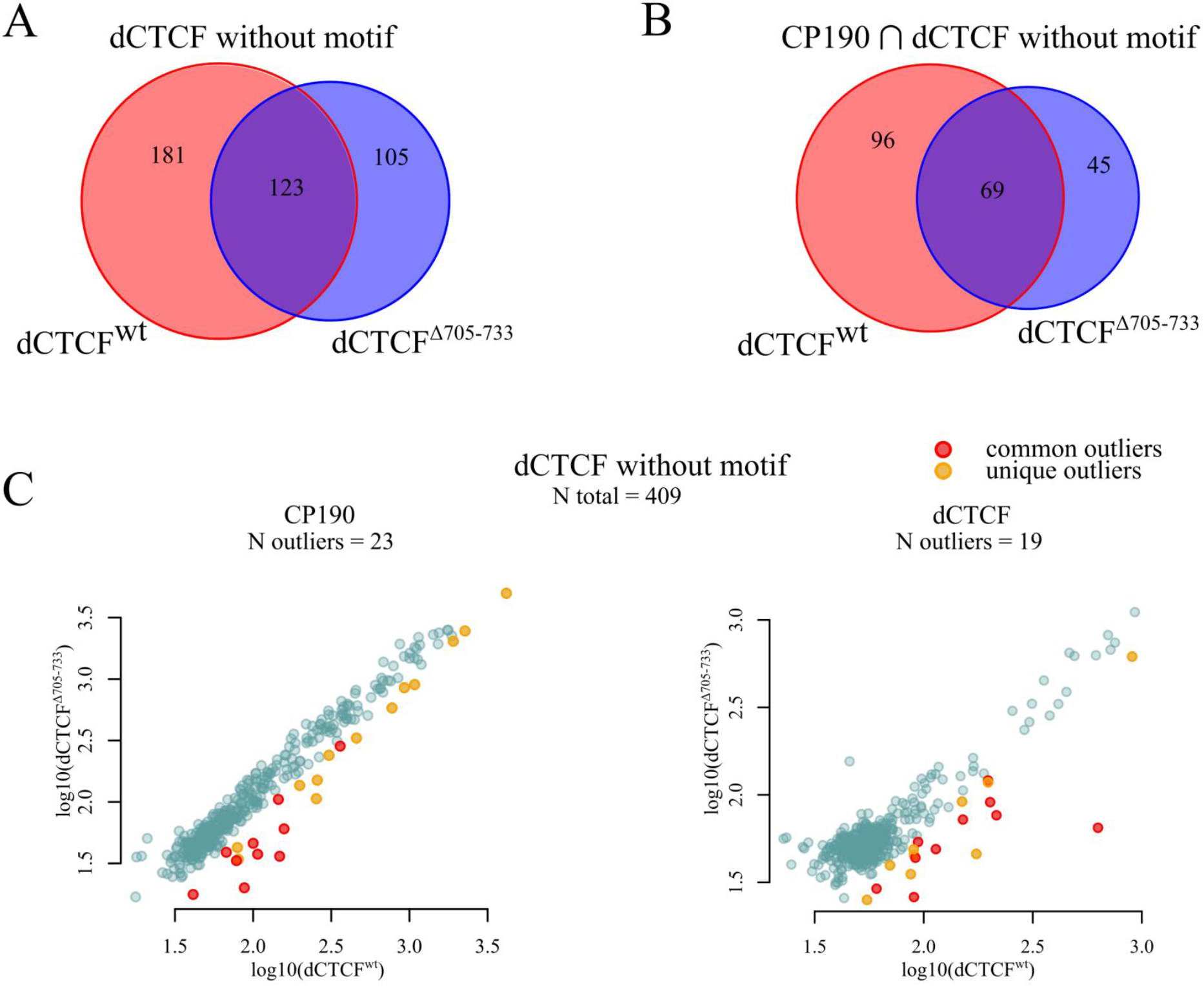
dCTCF and CP190 ChIP-Seq differences between *dCTCF*^*Δ705–733*^ and *dCTCF*^*wt*^ lines, considering only those dCTCF peaks without dCTCF motif sites. (A) Venn diagram showing the numbers of overlapping dCTCF peaks (without motif sites) between *dCTCF*^*Δ705–733*^ and *dCTCF*^*wt*^ lines. (B) Venn diagram showing the difference in the numbers of CP190 peaks overlapping with dCTCF (without motif sites) in *dCTCF*^*Δ705–733*^ and *dCTCF*^*wt*^ lines. (C) Comparison of the dCTCF peaks identified in the dCTCF and CP190 ChIP-Seq results (without motif sites, *dCTCF*^*Δ705–733*^ and *dCTCF*^*wt*^ merged) from *dCTCF*^*Δ705–733*^ and *dCTCF*^*wt*^ lines. Outliers were determined using linear regression (see Materials and Methods). Peaks that were determined to be outliers in both the CP190 and dCTCF signal analyses were denoted as “common outliers” (red). Peaks determined as outliers for only one of the proteins were denoted as “unique outliers” (yellow).

**Figure 7.**
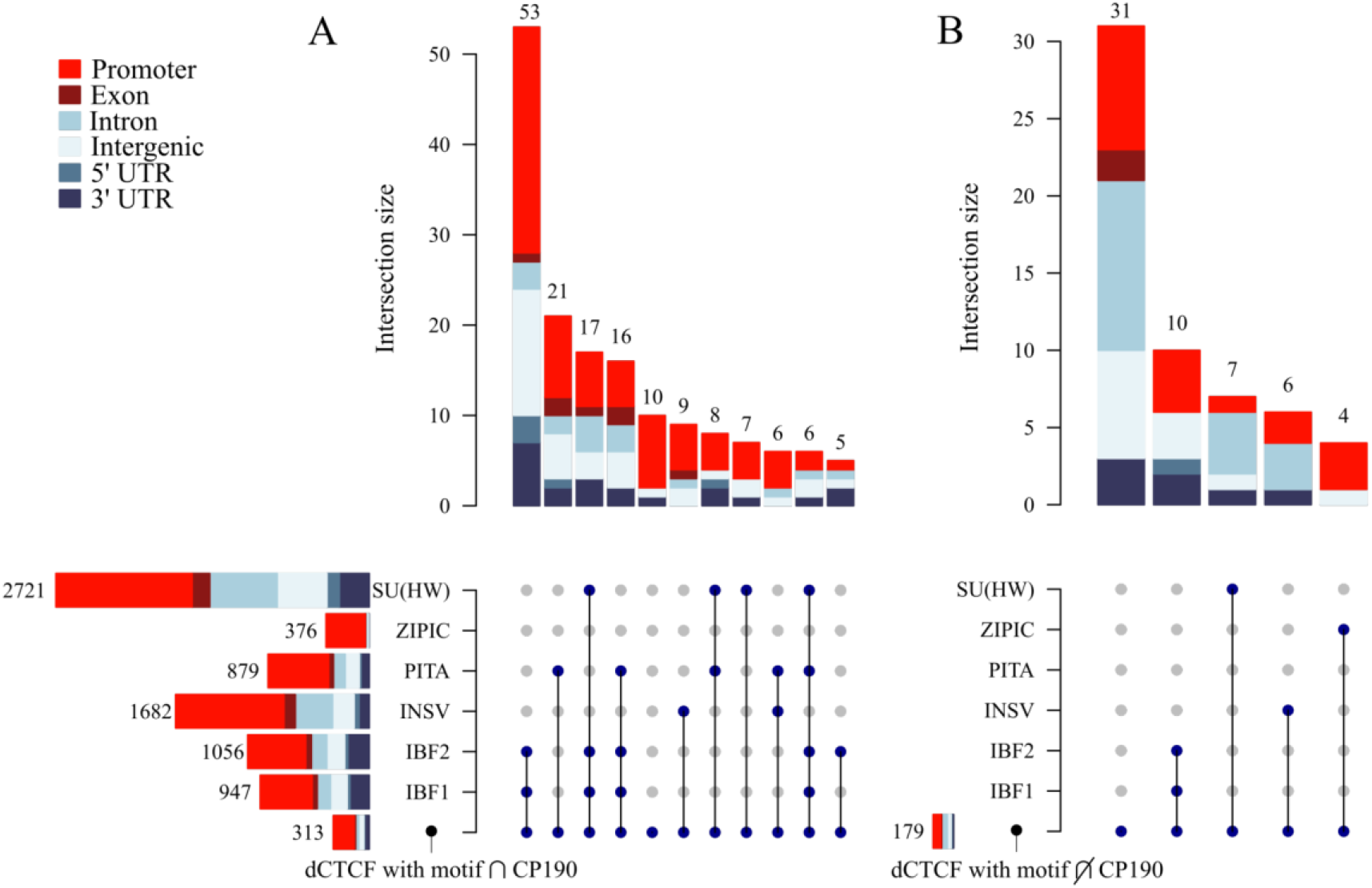
Colocalization of dCTCF peaks intersecting (A) and not intersecting (B) with CP190 with other proteins known by their interaction with CP190. Previously published data was used for the analysis (see Methods). Only peaks with a corresponding motif site were considered for each of the proteins. Intersections with less than three common peaks are not shown.

